# Host Age Structure Defines Interactions with Pathogens: Grandparent Effect under Collaboration and Virulent Mutualism under Competition

**DOI:** 10.1101/2022.08.16.504183

**Authors:** Carsten O.S. Portner, Edward G. Rong, Jared A. Ramirez, Yuri I Wolf, Angelique P. Bosse, Eugene V. Koonin, Nash D. Rochman

## Abstract

**Background:** Symbiotic relationships are ubiquitous in the biosphere. Inter-species symbiosis is impacted by intra-specific distinctions, in particular, those defined by the age structure of a population. Older individuals compete with younger individuals for resources despite being less likely to reproduce, diminishing the fitness of the population. Conversely, however, older individuals can support the reproduction of younger individuals, increasing the population fitness. Parasitic relationships are commonly age structured, typically, more adversely affecting older hosts.

**Results:** We employ mathematical modeling to explore the differential effects of collaborative or competitive host age structures on host-parasite relationships. A classical epidemiological compartment model is constructed with three disease states: susceptible, infected, and recovered. Each of these three states is partitioned into two compartments representing young, potentially reproductive, and old, post-reproductive, hosts, yielding 6 compartments in total. In order to describe competition and collaboration between old and young compartments, we model the reproductive success to depend on the fraction of young individuals in the population. Collaborative populations with relatively greater numbers of post-reproductive hosts enjoy greater reproductive success whereas in purely competitive populations, increase of the post-reproductive subpopulation reduces reproductive success. However, in competitive populations, virulent pathogens preferentially targeting old individuals can increase the population fitness.

**Conclusions:** We demonstrate that, in collaborative host populations, pathogens strictly impacting older, post-reproductive individuals can reduce population fitness even more than pathogens that directly impact younger, potentially reproductive individuals. In purely competitive populations, the reverse is observed, and we demonstrate that endemic, virulent pathogens can oxymoronically form a mutualistic relationship with the host, increasing the fitness of the host population. Applications to endangered species conservation and invasive species containment are discussed.

## Background

Despite having acquired a positive connotation when used informally, the term “symbiosis” may refer to any sustained inter- (but not intra-) species relationship(1). This relationship can be mutualistic (beneficial for all species involved), commensal (beneficial or neutral for all species), or parasitic (beneficial for some and deleterious for others) as well as obligate or facultative (beneficial but not required for survival). The diversity of inter-species symbioses is broadened still by intra-species diversity such that the nature of the symbiosis varies among individuals within each species. Pathogenic (that is, involved in strongly asymmetric, parasitic relationships(2)) human viruses provide well-characterized examples. For almost all pathogenic viruses, a period of host immunity follows infection, compartmentalizing the population into two groups, only one of which would engage in symbiosis with the virus(3). For common viruses that confer lifelong immunity following initial infection (or vaccination), these two groups can be often well approximated by age whereby young individuals who have never been exposed are susceptible to infection and older individuals, who were most likely exposed when they were themselves younger, are immune.

Indeed, the age structure of human populations plays an important role in forecasting the impact of novel pathogens because, even among apparently susceptible hosts, increased age may be a predictor of increased(4, 5) or decreased(5, 6) morbidity and mortality. Although post-reproductive lifespans appear to be rare in nature, even among mammals(7), senescence has been observed in all major branches of the tree of life, from animals(8, 9) and plants(10) to yeast(11) and bacteria(12). Senescent members are inherently costly to the population as they consume shared resources but reproduce with reduced (down to zero) efficiency. However, senescent individuals also can increase the overall growth rate of populations if they increase the efficiency with which other members reproduce. Such collaborative populations foster selection for post-reproductive lifespans, commonly referred to as “the grandmother hypothesis,” best demonstrated among humans(13–15) and orca whales(16).

In this work, we sought to explore the impact of host age-structure dynamics on the introduction of a pathogen with age-dependent virulence. We consider two diverse host populations: a competitive population modelled after rotifers(17, 18) (as well as an additional case modeled after solitary bees(19, 20) with qualitatively similar behavior addressed in the supplement) and a collaborative population modelled after humans. We demonstrate that, in competitive populations, a virulent pathogen can paradoxically increase the population growth rate. Conversely, in collaborative populations, the loss of a single post-reproductive member due to infection can result in a growth rate drop that exceeds that resulting from the loss of a single reproductive member.

## Results

### Model Construction

A classical epidemiological compartment model (Figure 1A) was constructed with three disease states: susceptible, infected, and recovered. Each of these three states is partitioned into two compartments representing young, potentially reproductive, and old, post-reproductive, hosts, yielding 6 compartments in total. All hosts are born susceptible (at rate *k_B_*) to infection; age into the post-reproductive compartment (at rate *k_A_*); and die from causes unrelated to infection (at rate *k_D_*). Susceptible hosts may become infected (proportional to rate *k_I_*) and recover or die due to infection (at rate *k_R_* or the age-specific rates *k_DYI_* or *k_DOI_* respectively). Recovered hosts can lose immunity and return to a susceptible compartment (at rate *k_L_*). For each population, the birth rate, death rate, and rate of aging from young to old compartments were fixed across all simulations (for a complete list of all parameters, see Table 1).

**Figure 1.**
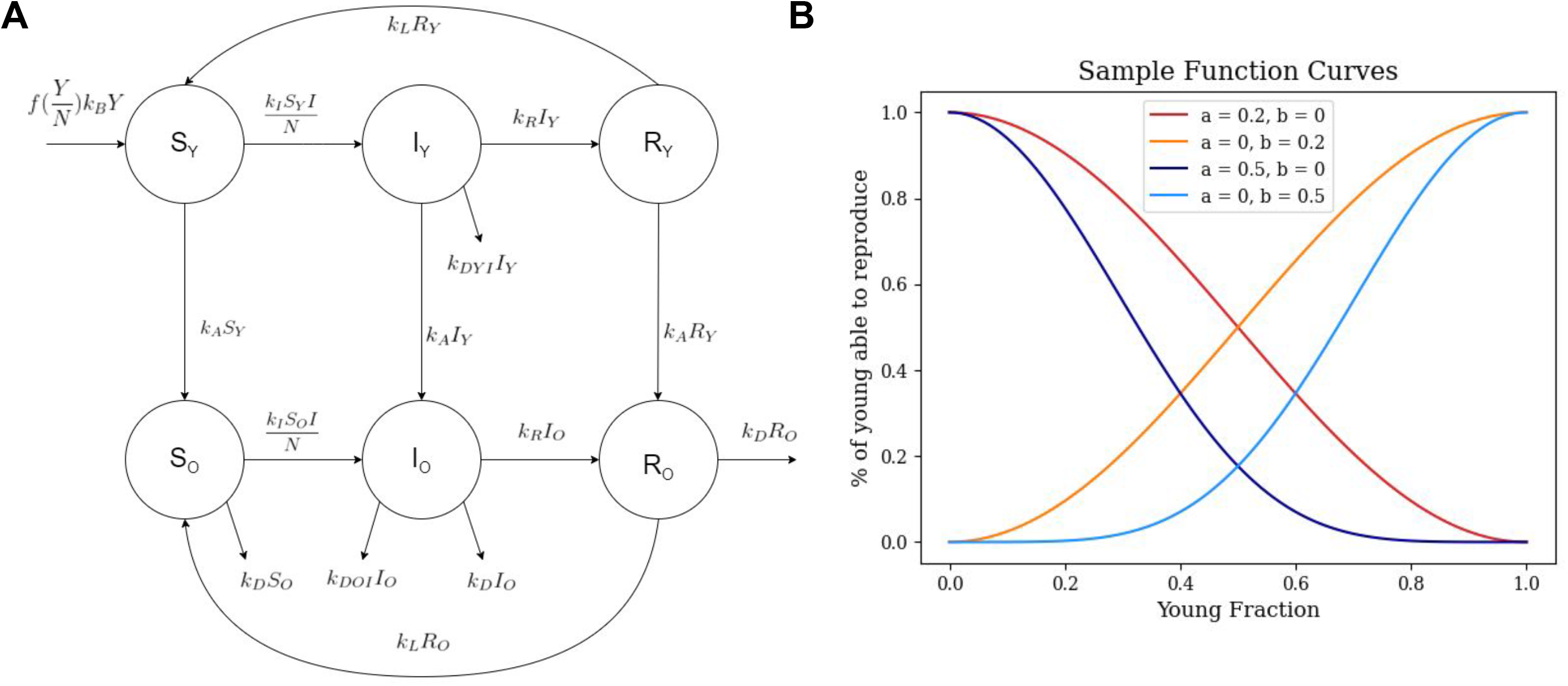
Model description. **A.** Compartment model diagram outlining three disease states each with two age compartments: one younger, potentially reproductive, and one post-reproductive. **B.** Functional dependence on the population age structure for the fraction of younger hosts which reproduce under collaborative, *a* = 0, 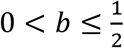, and competitive, *b* = 0, 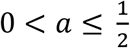, conditions.

**Table 1.**

|  | Human | Rotifer | Sweat Bee |
| --- | --- | --- | --- |
| Rate of Infection | 2 week<sup>-1</sup> | 4 days<sup>-1</sup> | 1.9 weeks<sup>-1</sup> |
| Rate of Recovery | 0.5 week<sup>-1</sup> | 0.059 days<sup>-1</sup> | 0.06 weeks<sup>-1</sup> |
| Rate of Loss of Immunity | 1 years<sup>-1</sup> | 0.059 days<sup>-1</sup> | 0.06 weeks<sup>-1</sup> |
| Birth Rate | 0.14 years<sup>-1</sup> | 1.3 days<sup>-1</sup> | 1.5 weeks<sup>-1</sup> |
| Aging Rate | 0.04 years<sup>-1</sup> | 0.125 days<sup>-1</sup> | 0.14 weeks<sup>-1</sup> |
| Post-Reproductive Death Rate | 0.022 years<sup>-1</sup> | 0.11 days<sup>-1</sup> | 0.14 weeks<sup>-1</sup> |
| Post-Reproductive Infection Death Rate | 0.25 years<sup>-1</sup> | 0.33 days<sup>-1</sup> | 1.4 weeks<sup>-1</sup> |

In order to model competition and collaboration between old and young compartments, we sought to make the reproductive success (which may represent the fraction of young hosts which are fertile or the probability that a birth will result in viable offspring) dependent on the proportion of the young individuals in the population, 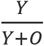, where *Y* and *O* are the cumulative sizes of all young and old compartments, respectively. First let us consider a collaborative population. In the limiting case where all but a single host (or mating pair) is post-reproductive, and aiding in the reproduction of that single host, we assume 100% reproductive success. The more collaborative the population, or conversely the greater the degree to which young hosts depend on post-reproductive hosts, the more steeply the reproductive success declines with relatively fewer post-reproductive hosts. Under conditions without any post-reproductive hosts, we assume the limiting case where the reproductive success is zero. We further assume this behavior saturates with respect to either limit.

The converse argument applies to competitive populations: reproductive success is inversely proportional to the relative number of post-reproductive hosts. Although cooperation and competition are not necessarily strictly symmetrical, for formal convenience, we considered the family of functions:

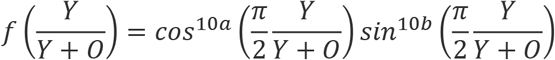

where *a* = 0, 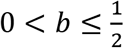 and *b* = 0, 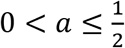 represent competitive and collaborative populations, respectively (Figure 1B), which possess the properties described above. We also emphasize that competition between parameter regimes is not assessed in this work. Only the relative fitness of disease free and endemic equilibria are considered.

Each simulation was conducted as follows. Parameters *a* and *b* were selected and the equilibrium age distribution, 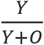, as well as the exponential growth rate, 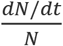 where *N* is the cumulative size of all compartments, were computed (for details, see Extended Data A). Compartments are initialized to respect the equilibrium age distribution, with infected compartments initialized to be 0.1% of the total population. The solution is then propagated using Python scipy method solve_ivp until a state of endemic equilibrium is reached where the measured rate of change of compartment proportions is negligible. The new growth rate at endemic equilibrium is then computed. For details, see Extended Data B.

### Loss of post-reproductive hosts is costly for collaborative populations

A human population was modelled consistent with an average reproductive life span of 25 years; post-reproductive life span of 45 years; a birth rate of 0.14 births/year; and reproductive success determined by *a* = {0.1,0.2,0.3,0.4,0.5}. The pathogen was modelled with an infectivity (the product of the rate of host contact and the probability of infection given contact) of 2/week, mean duration of infection 2 weeks, and mean duration of immunity 1 year for all hosts, but a probability of death roughly 1% for old hosts and zero for young hosts. These parameters well represent a range of human respiratory viruses(21) but we note this model does not account for the complex behavior associated with time varying contact rates(22, 23). The epidemic trajectory for *a* = 0.2 is shown in Figure 2A.

**Figure 2.**
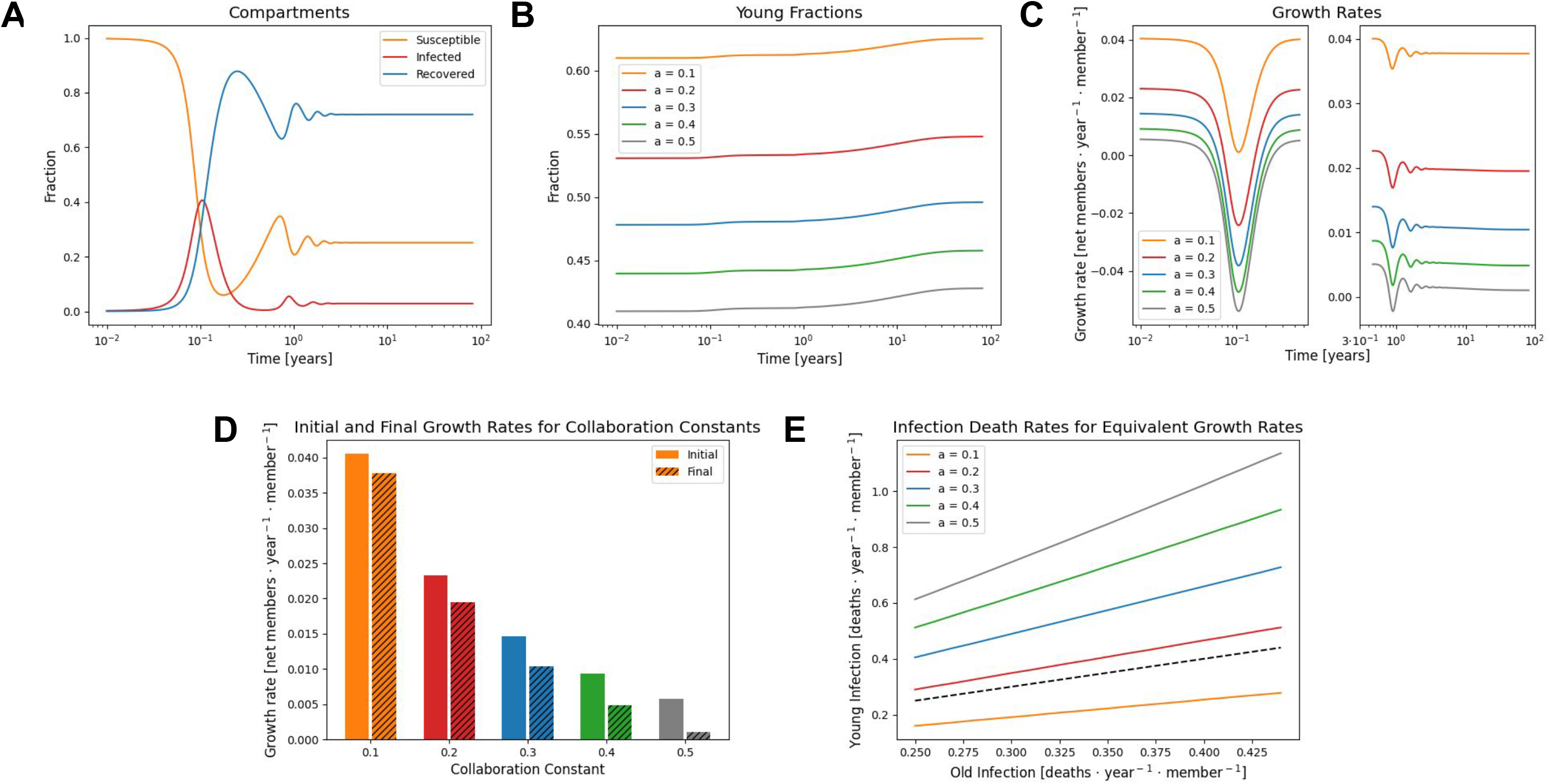
Value of post-reproductive hosts under collaboration. **A.** An example epidemic trajectory for *a* = 0.2 displaying the proportions of the three disease states. **B.** Epidemic trajectories for different degrees of collaboration with the younger, potentially reproductive, fraction of the population displayed. **C.** Population growth rate, 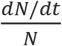, during the epidemic phase (left) and relaxation into endemicity (right). **D.** Comparison of initial, disease free, growth rates and final growth rates at endemic equilibrium. **E.** Equivalent infection death rates for younger, potentially reproductive, hosts resulting in the same growth rate suppression for variable infection death rates among post-reproductive hosts. For strongly collaborative populations the equivalent infection death rate for younger hosts is higher than for post-reproductive hosts.

Due to the loss of old hosts as a result of infection, the young fraction of the population increases throughout the epidemic phase and into endemicity (Figure 2B). In every case, the growth rate substantially declines (and can become negative) during the epidemic phase before gradually increasing (subject to oscillations) to a value reduced relative to the pre-epidemic level (Figure 2C). The cost to the population (decreased growth rate) as a result of endemic infection is greater for more collaborative populations (larger *a*, Figure 2D).

Death rate due to infection was then varied, ranging from 0.25/year to 0.45/year among old hosts, remaining at zero for young hosts. For each final growth rate, the equivalent death rate due to infection under conditions where only young hosts die as a result of infection was computed such that the final growth rate attained was the same for both models. Under sufficiently collaborative conditions, *a* ≥ 0.2, the equivalent death rate among young hosts resulting in the same growth rate reduction was greater than that among old hosts (Figure 2E). In other words, in a highly collaborative population, the cost of the loss of a post-reproductive member due to infection is greater than the cost of the loss of a young, potentially reproductive member.

The relative cost of losing young vs. post-reproductive hosts depends on the reproductive success function, 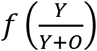. When reproductive success remains near one until 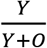 approaches one and then sharply declines, representing the case where the population is insensitive to the abundance of post-reproductive hosts, the loss of a young host always has a greater impact than the loss of a post-reproductive host (see Supplemental Figure 1).

### Virulent mutualism in competitive populations

A competitive population was modelled using the data available for rotifer populations, with an average reproductive life span of 8 days; post-reproductive life span of 9 days; a birth rate of 1.3/day(17); and reproductive success determined by *b* = {0.1,0.2,0.3,0.4,0.5}. The pathogen was modelled with an infectivity of 4/day, mean duration of infection 3 days, and mean duration of immunity 17 days for all hosts, but a probability of death near 100% for old hosts and zero for young hosts. These parameters, apart from age dependence, are modelled after a fungal pathogen of the genus *Rotiferophthora*(18). The epidemic trajectory for *b* = 0.2 is shown in Figure 3A. Note that the timescale of the epidemic is much shorter than that in the model of a collaborative population due to the major difference in member lifespans, and oscillations are not observed in these trajectories.

**Figure 3.**
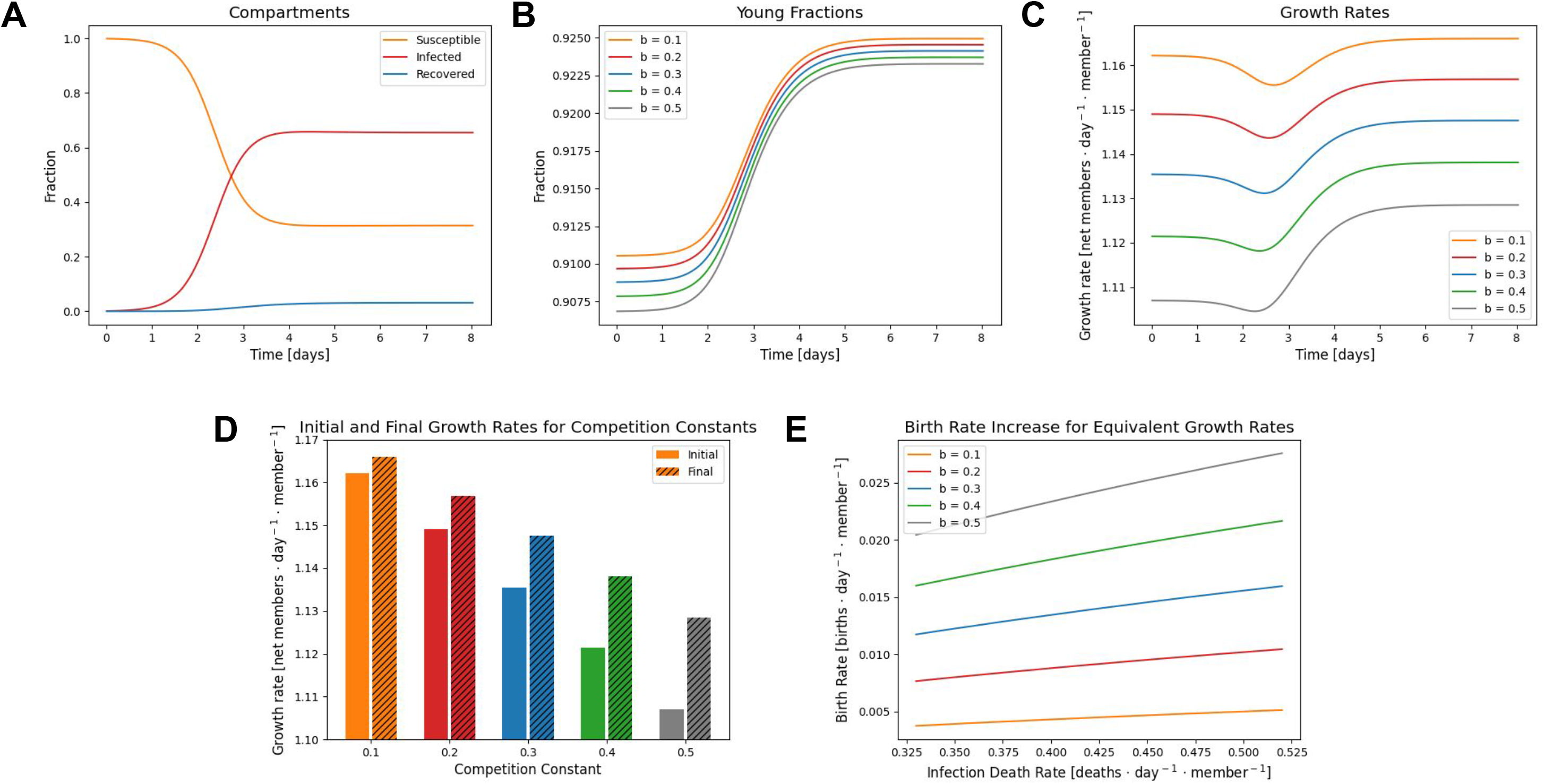
Emergence of virulent mutualists under competition. **A.** An example epidemic trajectory for *b* = 0.2 displaying the proportions of the three disease states. **B.** Epidemic trajectories for different degrees of competition with the younger, potentially reproductive, fraction of the population displayed. **C.** Population growth rate, 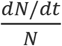, over the course of the epidemic which quickly reaches a state of endemic equilibrium. **D.** Comparison of initial, disease free, growth rates and final growth rates at endemic equilibrium. **E.** Equivalent increase in birth rate resulting in the same growth rate increase for variable infection death rates among post-reproductive hosts. For strongly competitive populations the equivalent birth rate increase can reach 2%.

Due to the loss of old hosts as a result of infection, the young fraction of the population increases logistically over the epidemic phase and into endemicity (Figure 3B). While the exponential growth rate (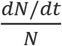, which is not the birth rate) still declines during the epidemic phase due to the loss of post-reproductive members, it remains positive and the endemic growth rate is elevated relative to the disease-free state (Figure 3C). The benefit to the population (increased growth rate) as a result of endemic infection is greater for more competitive populations (larger *b*, Figure 2D).

As for the collaborative case, the death rate due to infection was then varied, ranging from 0.33/day to 0.52/day among old hosts, remaining at zero for young hosts. For each final growth rate, the equivalent birth rate for a disease-free state was computed such that the final growth rate attained was the same for both models. Note that even a 1% change in the exponential growth rate under these conditions results in a 50% larger population within the time span of 3 generations. The introduction of a virulent pathogen significantly increased the population growth rate, in some cases equivalent to increasing the birth rate by more than 2% (Figure 3E). An additional competitive population was loosely modelled after sweat bees(19) introduced to a pathogen resembling acute bee paralysis virus(20), resulting in qualitatively similar behavior (see Supplemental Figure 2).

## Discussion

Parasites, by definition, decrease host fitness and the introduction of a parasite into one or more competing host species can substantially alter the ecological landscape resulting in or preventing the extinction of one competitor(24, 25). Symbionts that are parasitic in the absence of inter-species host competition but which increase the competitiveness of infected hosts (for example, through allelopathy(26)) establish a seemingly surprising, mutualistic relationship. Similarly, host competition can admit the persistence of a parasite under conditions where persistence within a single host population is not supported(25). Furthermore, even within a single host species, symbionts that are parasitic in one environment can provide a fitness advantage due to specialized adaptations in another environment, for example, promoting resistance to drought(27).

Parasitic interactions can also substantially alter intra-species competition due to variability in tolerance or transmissibility influenced by genomic or nongenomic factors including host density(28) and age. In previous human epidemics, advanced age has been commonly associated with increased(4, 5) or occasionally decreased(5, 6) morbidity and mortality. More generally, species lifespans are likely influenced by pathogen interactions. Populations of species with short lifespans are able to clear pathogens faster, and being short-lived decreases the probability of crossing the species barrier, which typically requires major adaptations to emerge within the first infected novel host to mediate efficient transmission(29).

Age structure substantially alters transmission dynamics, and can reduce the likelihood of an epidemic while increasing the magnitude in the event of an outbreak(30). The presence of immune post-reproductive hosts decreases the density of susceptible hosts and reduces the rate of infection; however, susceptible, and senescent, post-reproductive hosts can increase the cumulative number of mortalities. The disparity in predicted outcomes between homogenous (which may already lead to diverse results(31, 32)) and age-structured populations is exacerbated in the event of vaccination(33).

Here we demonstrate that host age structure, associated with differential susceptibility to a parasite, can determine whether a virulent pathogen, as defined by its relationship with individual hosts, is a parasite or a mutualist with respect to the population as a whole. In competitive populations where older, post-reproductive hosts share resources with younger, potentially reproductive hosts, virulent pathogens that disproportionately affect non-reproductive hosts increase the population growth rate. Conversely, in collaborative populations where older, post-reproductive hosts aid in the reproduction of younger hosts, the cost of the loss of a non-reproductive host can be even greater than that of a young host.

Although post-reproductive lifespans appear to be rare, even among mammals(7), most if not all cellular life forms exhibit some form of senescence(8–12). Therefore, although advanced social roles for post-reproductive members have not yet been identified outside of mammals, costs and potentially benefits associated with the presence of senescent hosts are likely widespread even for unicellular life. Among prokaryotes, horizontal gene transfer from dead, maximally senescent, cells can prevent irreversible genomic deterioration, Muller’s rachet(34). In addition to chronological age, altering the stationary cell cycle phase distribution(35, 36) can modify the fitness landscape by altering phase-dependent gene expression(37) potentially delegating different functions to individuals in different phases. Thus, collaborative populations might be far more common across the diversity of life than is immediately obvious.

These, often complex, impacts of age structure have to be taken into account prior to formulating a strategy for intervention against a pathogen, mitigation of invasive species, or preservation of an endangered population(38). For example, food provisioning both increases resource availability and host aggregation, potentially amplifying pathogen invasion(39). Thus, in the context of competition, which may itself be magnified as a result of host aggregation, the effect of a pathogen likely enhances conservation efforts, whereas in the case of collaboration, this straightforward intervention could lead to population decline.

## Supporting information

SupplementaryFigures_and_ExtendedData

## Author contributions

YIW, APB, NDR, and EVK designed the study; COSP, EGR, and JAR programed the model; COSP, EGR, JAR, YIW, and NDR conducted the analysis; COSP, NDR, and EVK wrote the manuscript that was edited and approved by all authors.

## Acknowledgements

The authors thank Koonin group members for helpful discussions. YIW, NDR, and EVK are supported by the Intramural Research Program of the National Institutes of Health.

## Supplemental Figure legends

**Supplemental Figure 1. Alternative model for collaboration among humans**

**A.** Functional dependence on the population age structure for the fraction of younger hosts which reproduce under collaborative, *a* = 0, 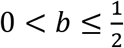, and competitive, *b* = 0, 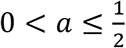, conditions. Alternative functional form from that used in the main text. **B.** An example epidemic trajectory for *a* = 0.2 displaying the proportions of the three disease states. **C.** Epidemic trajectories for different degrees of collaboration with the younger, potentially reproductive, fraction of the population displayed. **D.** Population growth rate, 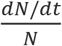, during the epidemic phase (left) and relaxation into endemicity (right). **E.** Comparison of initial, disease free, growth rates and final growth rates at endemic equilibrium. **F.** Equivalent infection death rates for younger, potentially reproductive, hosts resulting in the same growth rate suppression for variable infection death rates among post-reproductive hosts.

**Supplemental Figure 2. Competition among sweat bees**

**A.** An example epidemic trajectory for *b* = 0.2 displaying the proportions of the three disease states. **B.** Epidemic trajectories for different degrees of competition with the younger, potentially reproductive, fraction of the population displayed. **C.** Population growth rate, 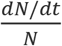, over the course of the epidemic which quickly reaches a state of endemic equilibrium. **D.** Comparison of initial, disease free, growth rates and final growth rates at endemic equilibrium. **E.** Equivalent increase in birth rate resulting in the same growth rate increase for variable infection death rates among post-reproductive hosts.

