## SupplementaryFigures_and_ExtendedData for "Host Age Structure Defines Interactions with Pathogens: Grandparent Effect under Collaboration and Virulent Mutualism under Competition"

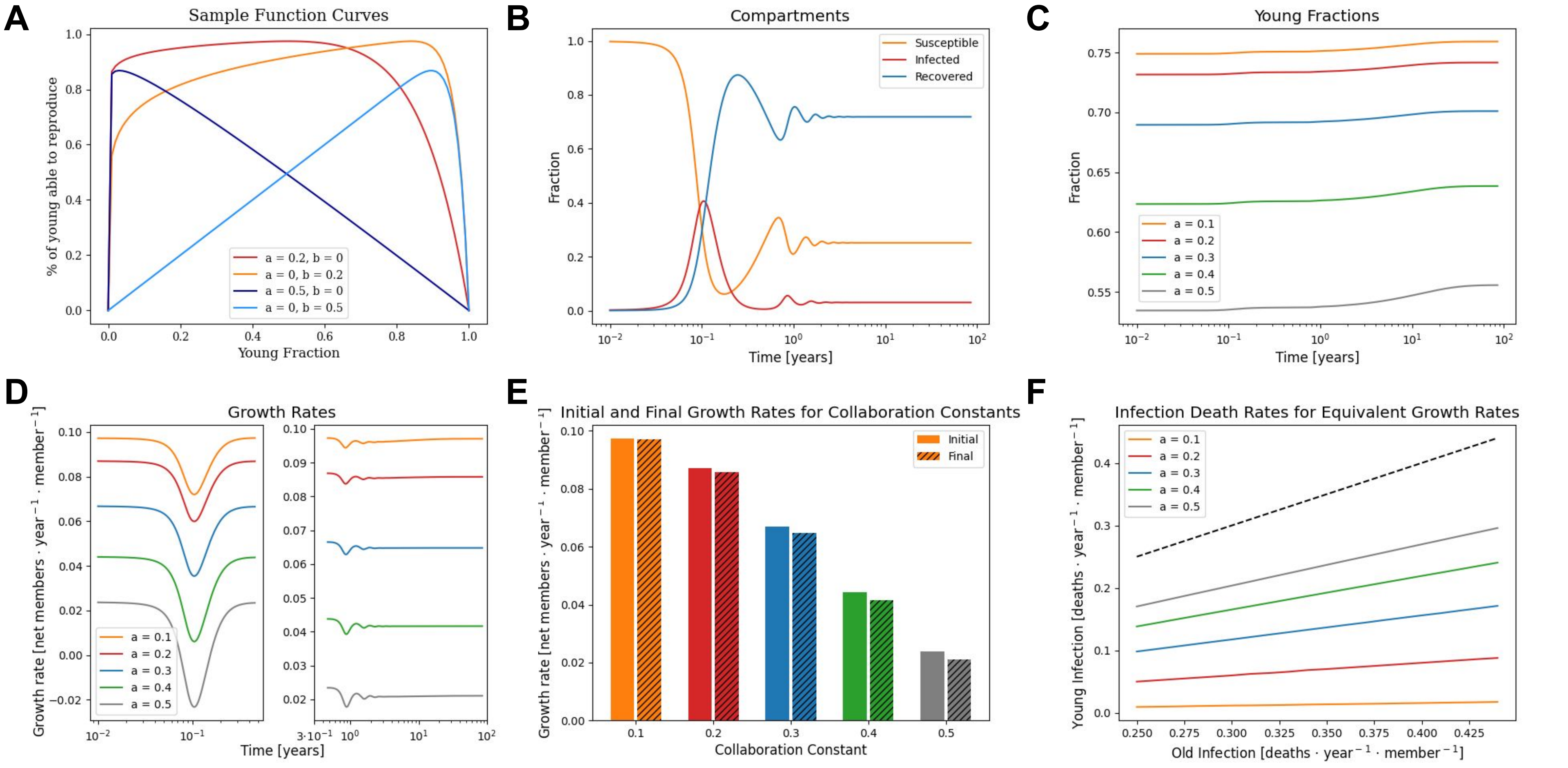

A
Sample Function Curves
% of young able to reproduce
Young Fraction
a = 0.2, b = 0
a = 0, b = 0.2
a = 0.5, b = 0
a = 0, b = 0.5
B
Compartments
Fraction
Time [years]
Susceptible
Infected
Recovered
C
Young Fractions
Fraction
Time [years]
a = 0.1
a = 0.2
a = 0.3
a = 0.4
a = 0.5
D
Growth Rates
Growth rate [net members · year⁻¹ · member⁻¹]
Time [years]
E
Initial and Final Growth Rates for Collaboration Constants
Growth rate [net members · year⁻¹ · member⁻¹]
Collaboration Constant
Initial
Final
F
Infection Death Rates for Equivalent Growth Rates
Young Infection [deaths · year⁻¹ · member⁻¹]
Old Infection [deaths · year⁻¹ · member⁻¹]

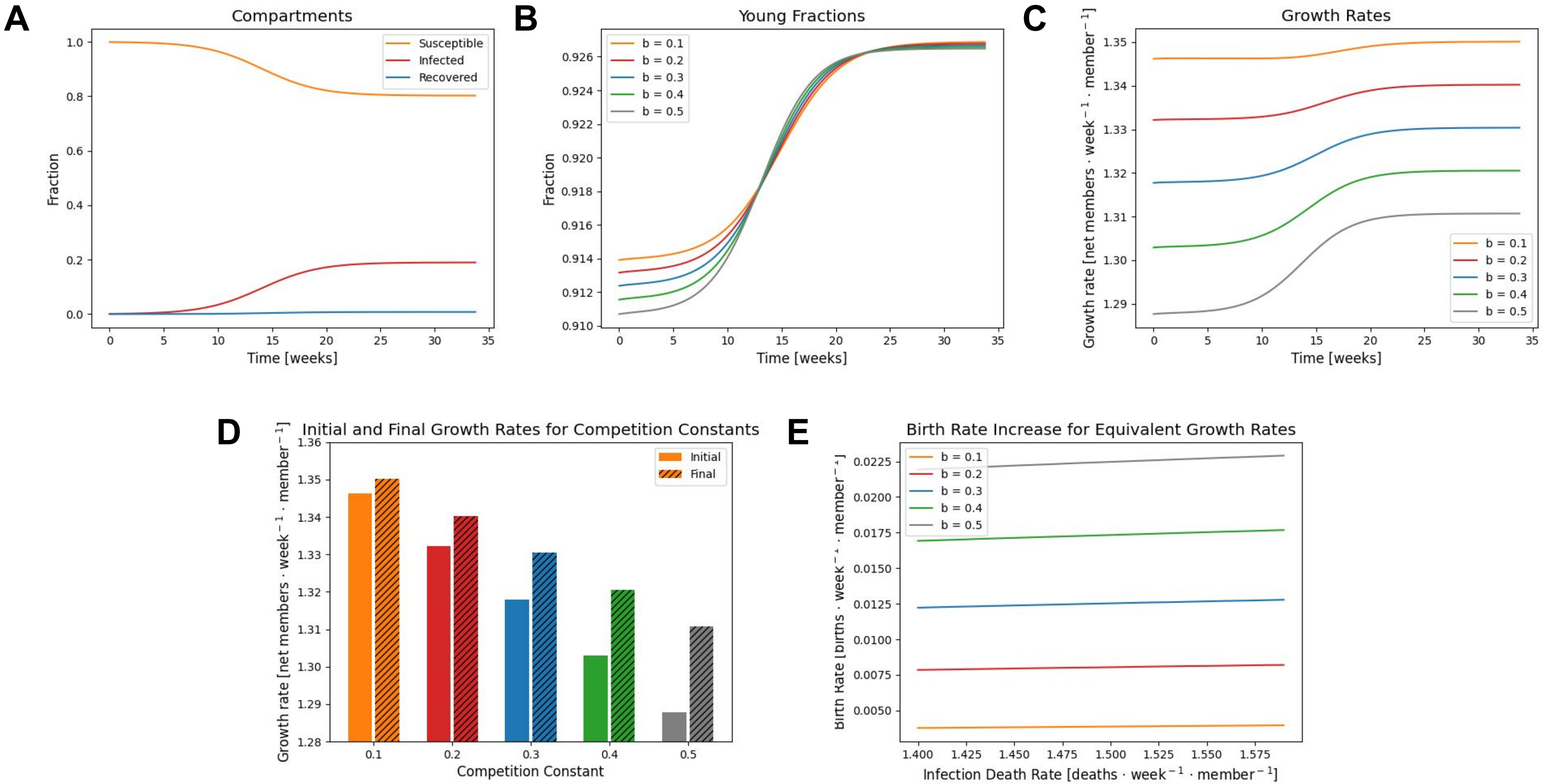

A
Compartments
Susceptible
Infected
Recovered
Fraction
Time [weeks]
B
Young Fractions
b = 0.1
b = 0.2
b = 0.3
b = 0.4
b = 0.5
Fraction
Time [weeks]
C
Growth Rates
Growth rate [net members · week⁻¹ · member⁻¹]
b = 0.1
b = 0.2
b = 0.3
b = 0.4
b = 0.5
Time [weeks]
D
Initial and Final Growth Rates for Competition Constants
Initial
Final
Growth rate [net members · week⁻¹ · member⁻¹]
Competition Constant
E
Birth Rate Increase for Equivalent Growth Rates
b = 0.1
b = 0.2
b = 0.3
b = 0.4
b = 0.5
Birth Rate [births · week⁻¹ · member⁻¹]
Infection Death Rate [deaths · week⁻¹ · member⁻¹]

**Extended Data A: Specification of age structure**

We begin with an age structured population, divided into two compartments of young, potentially reproductive members, $Y$, and older, post-reproductive members, $O$. The proportion of young hosts which are able to reproduce is goverend by the proportion of the population which is young:

$$f\left(\frac{Y}{Y+O}\right) = cos^{10a}\left(\frac{\pi}{2}\frac{Y}{Y+O}\right)cos^{10b}\left(\frac{\pi}{2}\frac{Y}{Y+O}\right)$$

unless otherwise specified, and:

$$f\left(\frac{Y}{Y+O}\right) = \left(1-\frac{Y}{Y+O}^{1000\left(\frac{1}{2}-a\right)}\right)\frac{Y}{Y+O}^{1000\left(b-\frac{1}{2}\right)}$$

for Supplementary Figure 1 where $a=0$, $0<b<\frac{1}{2}$ or $b=0$, $0<a<\frac{1}{2}$. Reproductive hosts give birth at rate $k_B$ and age into the post-reproductive compartment at rate $k_A$. Post-reproductive hosts die at rate $k_D$. These dynamics are represented by the system of equations:

$$\begin{bmatrix} Y \\ O \end{bmatrix}' = \begin{bmatrix} f\left(\frac{Y}{Y+O}\right)k_B - k_A & 0 \\ k_A & -k_D \end{bmatrix}\begin{bmatrix} Y \\ O \end{bmatrix}$$

This system may be utilized to derive the steady state age distribution, defined by the ratio $\frac{Y}{Y+O}$:

$$0 = \left(\frac{Y}{Y+O}\right)' = \frac{Y'}{(Y+O)} - \frac{Y(Y+O)'}{(Y+O)^2} = \frac{Y}{Y+O}\left(f\left(\frac{Y}{Y+O}\right)k_B + k_D - k_A\right) - \left(\frac{Y}{Y+O}\right)^2\left(f\left(\frac{Y}{Y+O}\right)k_B + k_D\right)$$

which may be solved numerically for arbitrary $f\left(\frac{Y}{Y+O}\right)$.

**Extended Data B: Construction of epidemiological compartment model**

We may now introduce the age structured population defined in the previous section, initialized with the steady state age distribution, to a pathogen. Hosts may be susceptible ($S_Y$,$S_O$), infected and infectious ($I_Y$,$I_O$), or recovered and immune ($R_Y$,$R_O$). Susceptible young/older hosts are infected at rate $k_I\frac{I}{N}S_{Y/O}$ where $I \equiv I_Y + I_O$ and $N$ is the sum over all compartments, the total population. The death rate of young and old infected hosts due to infection may differ, $k_{DYI/DOI}$, and all recovered hosts may return to a susceptible compartment at rate $k_L$ due to a loss of immunity. These dynamics yield the system of equations:

$$d\begin{bmatrix} S_Y \\ S_O \\ I_Y \\ I_O \\ R_Y \\ R_O \end{bmatrix}/dt = \begin{bmatrix} f(\frac{Y}{N})k_B - \frac{k_I I}{N} - k_A & 0 & f(\frac{Y}{N})k_B & 0 & f(\frac{Y}{N})k_B + k_L & 0 \\ k_A & -\frac{k_I I}{N} - k_D & 0 & 0 & 0 & k_L \\ \frac{k_I I}{N} & 0 & -k_R - k_{DYI} - k_A & 0 & 0 & 0 \\ 0 & \frac{k_I I}{N} & k_A & -k_R - k_{DOI} - k_D & 0 & 0 \\ 0 & 0 & k_R & 0 & -k_L - k_A & 0 \\ 0 & 0 & 0 & k_R & k_A & -k_L - k_D \end{bmatrix}\begin{bmatrix} S_Y \\ S_O \\ I_Y \\ I_O \\ R_Y \\ R_O \end{bmatrix}$$

Each simulation is conducted as follows. Parameters $a$ and $b$ are selected and the equilibrium age distribution, $\frac{Y}{Y+O}$, as well as the growth rate, $(dN/dt)/N$ where $N$ is the cumulative size of all compartments, are computed. Compartments are initialized to respect the equilibrium age distribution with infected compartments initialized to be 0.1% of the total population. The solution is then propagated using Python scipy method solve_ivp until a state of endemic equilibrium is reached where the measured rate of change of compartment proportions is negligible:

$$|\frac{d\frac{Y}{N}}{dt}|\frac{N}{Y} + |\frac{d\frac{O}{N}}{dt}|\frac{N}{O} + |\frac{d\frac{S}{N}}{dt}|\frac{N}{S} + |\frac{d\frac{I}{N}}{dt}|\frac{N}{I} + |\frac{d\frac{R}{N}}{dt}|\frac{N}{R} < 0.0001$$
